# A large-scale study on the nocturnal behavior of African ungulates in zoos and its influencing factor

**DOI:** 10.1101/2023.06.13.544771

**Authors:** Jennifer Gübert, Max Hahn-Klimroth, Paul W. Dierkes

## Abstract

This study analyzed the nocturnal behavior of 196 individuals of 19 ungulate species in 20 zoos in Germany and the Netherlands. To the best of our knowledge, this is the first description of nocturnal behavior for some of the species. The importance of a wide range of possible factors influencing nocturnal behavior is discussed. Specifically, the behavioral states of standing and lying were analyzed, evaluating the proportion and number of phases in each behavior. The underlying data consists of 101,629 hours of video material from 9,239 nights. BOVIDS, a deep learning-based software package, was used to analyze the recordings. The analysis of the influencing factors was based on a random forest regression and a SHAP analysis. The results indicate that age, body size and feeding type are the most important factors influencing nocturnal behavior across all species. There are strong differences between the zebra species and the observed Cetartiodactyla as well as White Rhinos. The main difference is that zebras spend significantly less time in a lying position than Cetartiodactyla.

## 1 Introduction

There are over 250 species of odd-toed and even-toed ungulates, and Africa in particular is home to around 100 representatives of these orders (Ezenwa et al., 2006; IUCN, 2022). Ungulates are moreover characteristic animals of the African landscape, and also include some of the most widely kept zoo animals (Rose and Robert, 2013), as well as the most prominent farm animals such as cattle, sheep, pigs, or horses (FAO, 2013). As the African continent consists of multiple habitats, including different climatic and vegetation zones, African ungulates already show a great diversity in body measurements or physiology.

Behavioral patterns of large herbivores are known to vary greatly between day and night, while most ungulates show a diurnal to crepuscular cycle which conditions that they are most active during the day (Bennie et al., 2014; Gravett et al., 2017; Wu et al., 2018). For wild Arabian oryx it is known that especially during the colder months, sleep behavior shifts heavily into the night (Davimes et al., 2018). Nevertheless, for many ungulate species, only very few studies on the nocturnal behavior are available, both in the wild and in zoos (Berger, 2010; Rose and Robert, 2013). In particular, the sleeping behavior of many ungulates is poorly studied (Lyamin et al., 2021). On the other hand, the nocturnal behavior of very prominent ungulates like giraffes (Grzimek, 1956; Tobler and Schwierin, 1996; Seeber et al., 2012; Sicks, 2016; Burger et al., 2021), elephants (Gravett et al., 2017), or farm animals like cattle (Ruckebusch, 1972; Ternman et al., 2014; Fukasawa et al., 2018) is well studied. However, there are, to the best of our knowledge, many other ungulates whose nocturnal behavior has not been analyzed currently.

The knowledge about species’ behavior is not only scientifically interesting but it is also crucial for successful animal management and husbandry (Rose and Riley, 2021). This can lead to improved holding conditions and thus can benefit animal welfare (Berger, 2010; Brando and Buchanan-Smith, 2018; Walsh et al., 2019; Rose and Riley, 2021). Early detection of conspicuous behavioral patterns may indicate reduced animal welfare due to disease or stress, and zoos could quickly take countermeasures by measuring these behavioral patterns. For instance, sleep is an important factor of well-being (Hänninen et al., 2008; Fukasawa et al., 2018; Northeast et al., 2020). In particular, the monitoring of REM sleep allows us to draw conclusions about stress in giraffes (Sicks, 2016). Moreover, it is known to be a prognostic indicator during infectious disease in rabbits (Toth et al., 1993). Knowledge about a species’ expected behavior is required for those monitoring tasks. However, observing nocturnal behavior is challenging, especially in the wild. The analysis of animal’s behavior in zoos is a helpful tool to generate a broader knowledge and can be used as a good basis and starting information for observations in the field (Kleiman, 1992; Ryder and Feistner, 1995). Studies conducted in zoos have some advantages because they provide consistent and better access to animals and easier conditions for data collection (Ryder and Feistner, 1995), which is critical for large-scale studies. Of course, zoo animals have a quite different ecology from wild living conspecifics. Therefore, it is possible that zoo animals show different activity budgets. However, many characteristics of zoo animals and their wild conspecifics remain the same such that observations in zoos help us to learn about the species’ behavior in the wild (Rees, 2023).

In this publication, a large-scale study is conducted to contribute to the knowledge of the nocturnal behavior of African ungulates. The behavioral poses “standing”, “lying – head up” (LHU) and “lying – head down” (LHD) were analyzed. LHD is the typical REM (rapid eye movement) sleep posture and can be used to estimate REM sleep. The large advantage in contrast to physiological measurements is that REM sleep can be estimated by completely non-invasive methods on video recordings. Although, REM sleep can only be approximated by its typical body posture, this approximation is known to be decent, in particular for longer periods of LHD (Ternman et al., 2014; Lyamin et al., 2021; El Allali et al., 2022).

The current study examined the nocturnal behavior of 19 species of the orders Perissodactyla and Cetartiodactyla in 20 zoos. A total of 9,239 nights of video recordings from 196 animals are analyzed. This dataset allows us to analyze the importance of different factors influencing nocturnal behavior using random forest regression followed by SHAP analysis. The extensive amount of data could be analyzed using BOVIDS (Behavioral Observations by Videos and Images using Deep-Learning Software), a deep-learning software package for pose estimation that was developed specifically for this task (Gübert et al., 2022). Pose estimation is the machine learning task to automatically identify a pose, like standing or lying, from given image material. For some of the species studied, the current study provides the first description of nocturnal behavior that we know of and greatly improves the available data for other species.

## 2 Material and Methods

### 2.1 Ethogram

This study concentrates on a basic description of behaviors which can be detected via pose estimation. Thus, the main behaviors “standing” and “lying” are distinguished. Lying is furthermore split up into “lying -head up (LHU)” and “lying -head down (LHD)”. If an animal is not present or the behavior cannot be inferred from the recording, the category “out” is given. The investigated behaviors are defined in the following ethogram (Table 1 and Figure 1) based on Gübert et al. (2022).

**Table 1:** Ethogram for the studied ungulates, including the description of the three observed behavioral poses.

| Behavioral pose |  | Definition |
| --- | --- | --- |
| Standing | Standing | The animal stands in an upright position. Other behaviors like feeding, resting, walking, or ruminating can occur simultaneously. |
| Lying | Lying – head up (LHU) | The animal is in a sternal recumbency with the trunk touching the ground. Its head is lifted. |
|  | Lying – head down (LHD) | Cetartiodactyla: The animal is in a sternal recumbency with the trunk touching the ground (like in Lying - head up) but its head is resting on the ground.<br>Perissodactyla: The animal is lying in a lateral recumbency. |

**Figure 1:**
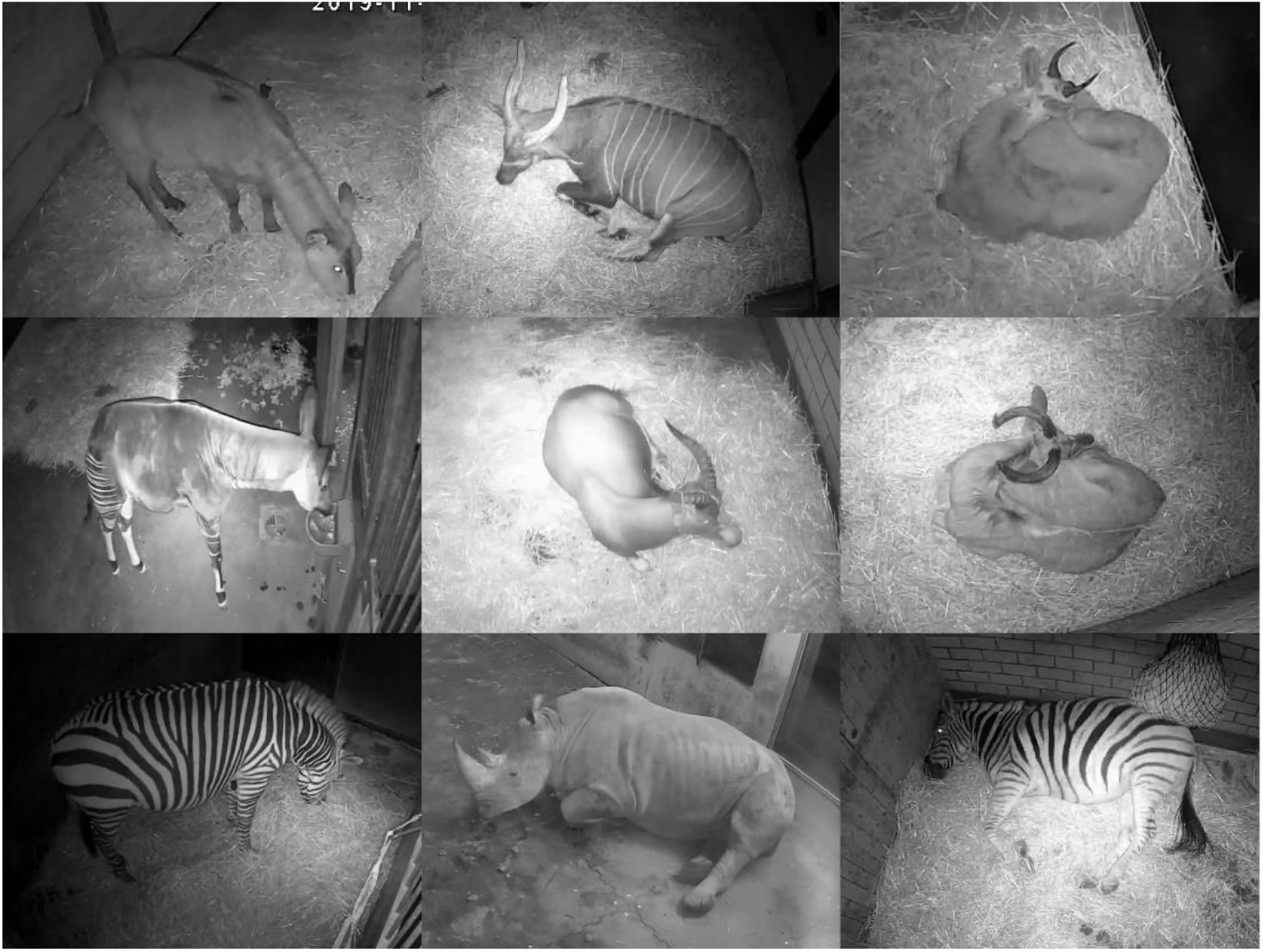
The three observed behavioral poses: standing (first column) and lying (second and third column). Lying is distinguished further into lying -head up (second column) and lying -head down (third column). The postures are shown by different species: Waterbuck, Bongo, Mountain Reedbuck (first row), Okapi, Blesbok, Greater Kudu (second row), Mountain Zebra, White Rhino, and Plains Zebra (third row).

LHD is the typical REM sleep posture and thus can be used to estimate REM sleep. This is based on the fact that due to postural atonia the animal’s head needs to be rested in REM sleep (Lima et al., 2005; Zepelin et al., 2005). It is a common method to estimate REM sleep by the LHD position, recent studies using this method include studies of Cetartiodactyla, like Common Eland (Zizkova et al., 2013), Giraffe (Seeber et al., 2012), Dromedary Camel (El Allali et al., 2022), and cattle (Ternman et al., 2014). Moreover, LHD is also used to estimate REM sleep in various studies on Perissodactyla’s behavior (Houpt, 1980; Pedersen et al., 2004; Greening and McBride, 2022).

### 2.2 Data collection

The data on which this study is based was collected by video recordings during the colder seasons (September to May) between 2017 and 2021 in 20 EAZA zoos in Germany (Zoologische Gärten Berlin (Tierpark and Zoo), Zoo Vivarium Darmstadt, Zoo Dortmund, Zoo Duisburg, Zoo Frankfurt, Zoom Erlebniswelt Gelsenkirchen, Erlebnis-Zoo Hannover, Zoo Heidelberg, Kölner Zoo, Zoo Krefeld, Opel-Zoo Kronberg, Zoo Landau in der Pfalz, Zoo Leipzig, Allwetterzoo Münster, Zoo Neuwied, Zoo Osnabrück, Zoologischer Garten Schwerin, Der Grüne Zoo Wuppertal) and the Netherlands (Königlicher Burgers Zoo Arnheim).

Two different datasets were used in the study. The larger dataset consisted of all evaluated data, i.e., 9,239 nights with a total of 101,629 hours. On this dataset the behavioral poses standing and lying were distinguished. On a subset of this dataset consisting of 6,265 nights with a total of 68,915 hours, lying was distinguished into LHU and LHD. Further details of the dataset can be found in the supplementary material (S1 Table).

The recordings on the complete dataset include 196 individuals and 19 ungulate species from the orders Perissodactyla and Cetartiodactyla. In this study the Perissodactyla include the species Plains Zebra (*Equus quagga*), Grevy’s Zebra (*Equus grevyi*) and Mountain Zebra (*Equus zebra*) of the family Equidae as well as the White Rhino (*Ceratotherium simum*) of the family Rhinocerotidae. Furthermore, the order Cetartiodactyla comprises the family Giraffidae, including Okapi (*Okapia johnstoni*), and the family Bovidae. African bovids studied are Greater Kudu (*Tragelaphus strepsiceros*), Sitatunga (*Tragelaphus spekii*), Bongo (*Tragelaphus eurycerus*), Common Eland (*Tragelaphus oryx*), Blesbok (*Damaliscus pygargus*), Common Wildebeest (*Connochaetes taurinus*), Roan Antelope (*Hippotragus equinus*), Sable Antelope (*Hippotragus niger*), Scimitar-horned Oryx (*Oryx dammah*), Addax (*Addax nasomaculatus*), Waterbuck (*Kobus ellipsiprymnus*), Mountain Reedbuck (*Redunca fulvorufula*) and African Buffalo (*Syncerus caffer*), whereby within the last species the two subspecies African Savannah Buffalo (*Syncerus caffer caffer*) and African Forest Buffalo (*Syncerus caffer nanus*) are distinguished due to morphological differences in body measurements. Finally, the data also contains recordings of the Arabian Oryx (*Oryx leucoryx*).

The video recordings were conducted by cameras capable of night vision due to integrated infrared emitters (Lupus LE139HD or Lupus LE338HD with the recording device LUPUSTEC LE800HD or TECHNAXX PRO HD 720P). The resolution of the recordings ranged from 704×576 px to 1920×1080 px and the framerate was 1 fps. The chosen recording time is from 7 pm to 6 am, so that no keepers were present during this time and all animals were already in the stable for some time at the beginning of the recording. Food was provided ad libitum. Furthermore, the natural light conditions which are known to be an important zeitgeber are comparable throughout the recording period (Merrow et al., 2005). One recording from 7 pm to 6 am is called a “night” and recordings on which an animal is not present for at least 20% of the time were dismissed. Additionally, to study quantities relating to the number of phases of a behavior shown during one night, nights with at least three occurrences of “out of view” were also dismissed. Ways of evaluating the video material are explained in the next section “Software”. The data was recorded continuously, so there is an exact time span for each behavioral sequence (Martin and Bateson, 2015).

### 2.3 Software

To evaluate the video recordings the software package BOVIDS was used (Hahn-Klimroth et al., 2021; Gübert et al., 2022). As this is a deep learning-based approach, testing data needs to be produced manually. At least two nights per individual were evaluated manually with the open-source software BORIS (Behavioral Observation Research Interactive Software), Version 7.7.3 (Friard and Gamba, 2016). In total, 517 nights corresponding to 5,687 hours of video material were annotated manually. This manually annotated data was the testing set. On this testing set, average f-scores of 0.992 ± 0.003 (lying), 0.979 ± 0.006 (standing) and 0.956 ± 0.006 (LHD) are achieved. Detailed information on the performance is given in the supplementary material (S2 Table and S3 Figure). The output of BOVIDS is a continuous sequence such that every 7 seconds of a recording were assigned a behavioral class. To minimize errors due to automated analysis, BOVIDS applies a set of so-called post-processing rules that dismiss short events. According to Gübert et al. (2022), phases of lying and standing below 5 minutes are dismissed as well as phases of LHD below 35 seconds.

For building and analyzing data scientific models Python’s scikit-learn library (Pedregosa et al., 2011) and the SHAP library (Lundberg and Lee, 2017) were used. Furthermore, data preparation was conducted by the Python library pandas (McKinney, 2010; Pandas Team, 2022). Regressions were done in R (R Core Team, 2014) and visualizations were prepared with ggplot2 (Wickham, 2016) or matplotlib (Hunter, 2007). For all figures and tables, a scientific species names’ abbreviation was defined. It consists of the first letter of the genus followed by the first three letters of the specific epithet. In all tasks an explained variance score, respectively R^2^, was considered moderate if it exceeds 0.13 and strong if it exceeds 0.26 (Cohen, 1988). Furthermore, the abbreviations SD, SEM and QD were used for the standard deviation, the standard error of the mean and the quartile deviation.

### 2.4 Data analysis

The behavioral poses were analyzed with respect to the proportion per night and the number of behavioral phases. Besides the description of the nocturnal behavior, the main goal of the study was to identify relevant influencing factors, so-called features, that notably influence the animal’s nocturnal behavior. Note that “proportion of lying” implies “proportion of standing” and that the “number of phases lying” estimates the “number of phases standing” up to ± 1 and thus it suffices to study the dependent variables “proportion lying”, “proportion LHD”, “number of phases lying”, and “number of phases LHD”. Indeed, as every lying phase is followed by a standing phase and vice versa, the number of phases lying is equal to the number of phases standing in the case standing-lying-standing-lying, overestimates the number of phases standing by one in the case lying-standing-lying and underestimates the number of phases standing by one in the case standing-lying-standing.

The analyzed features per night are the month of the recording (“month”), the species (“species”) of the observed individual, its sex (“sex”) and its age group (“age”) encoded as juvenile, subadult, or adult. The boundaries between these age categories are formed by the time of weaning and the sexual maturity of the species and are gathered from different sources (Grzimek, 2000; Puschmann et al., 2009; Groves and Leslie Jr, 2011; Rubenstein, 2011; Skinner and Mitchell, 2011; Tacutu et al., 2013; Myers et al., 2021). The according values are given in the supplementary material (S4 Table).

Furthermore, the indicator whether multiple species share a stable building or if the building only hosts one species is called “stable”. Moreover, the keeping zoo (“zoo”) was recorded. This variable is highly dependent on multiple non-measured features of the housing and husbandry like the feeding routine, the time being on an outdoor enclosure during day, the husbandry form like multi-species exhibits, etc. Also, “zoo” is statistically dependent on non-measured conditions independent from housing like weather.

The feature “species” implies a variety of other features. “Genus” and “family” are added as taxonomic features. “Food” corresponds to the dietary type: grazer, browser, or intermediate. Next, “species” contains information about the individual’s natural habitat (“habitat”) ranging from open to closed environments (IUCN, 2022). Last, “species” implies the average body measurements, in particular, the shoulder height “SH”, and the species’ average weight were used. All used values are given in detail in the supplementary material (S4 Table). The habitat is given by a number between 0 (open environments) and 3 (closed environments) following the categorization of the IUCN Red List of Threatened Species (IUCN, 2022). The species’ average body measurements and the dietary type are gathered from different sources (Grzimek, 2000; Puschmann et al., 2009; Groves and Leslie Jr, 2011; Rubenstein, 2011; Skinner and Mitchell, 2011; Tacutu et al., 2013; Myers et al., 2021).

The underlying dataset is high dimensional and contains numerical and categorical features. Furthermore, there is a large multicollinearity of the features and there are combinations of features that do not contain any sample. While dealing with such complicated datasets was almost impossible for a long time, increasing computational power and better understanding of various data scientific concepts made a more data driven approach find its way into life science studies (Boulesteix et al., 2012). For the dataset at hand a random forest regression was used because it has very few assumptions on the underlying distributions of the dataset. Further, it is robust to noisy measurements and outliers which is an important property as the raw data is annotated by means of a deep learning-based software package. Finally, the random forest regression allows to derive measures of feature importance which are a natural indicator of the influence of single features (Fraser, 1965; Parveen et al., 2013; Beraha et al., 2019). It is important to note that features that contribute significantly to the result in classical statistical tests are also considered relevant in a random forest regression and vice versa (Chicco and Jurman, 2021; Saarela and Jauhiainen, 2021).

Random forest regression is a predictive supervised learning model. Thus, it computes an estimate of the dependent variable based on the features. A random forest consists of several independent random decision trees. To build one decision tree, a bootstrap sample from the given data is drawn. At every node of the decision tree, one feature is selected among a random collection of features and the dataset is split optimally into two disjoint subsets based on this feature’s value. Among the random collection of features, the feature with the most optimal partition of the dataset is chosen. This splitting procedure continues recursively on the two subsets. In the contribution at hand, the random forest regressor consists of 300 decision trees each. The minimum number of samples per leaf and per split are set to 120 and the maximum depth of a decision tree is four to prevent overfitting. The maximum number of randomly selected features is set to 66% of the total number of features which is known to work well on smaller datasets at the cost of higher computational time. The split’s performance is measured with respect to Friedman’s MSE. The goodness of fit of the random forest regressor is assessed by the explained variance R^2^ between the model’s prediction and the original data because the explained variance is known to work decently with tree classifiers (Chen et al., 2020). Given the dataset R^2^ is defined as the corresponding mean of its bootstrap estimate (Efron, 1979; Efron, 1981). In the modelling part, balancing the influence that a single individual has on the model’s properties is necessary. Thus, for the random forest regression, equally many nights (n = 50) are sampled uniformly at random per individual. If more nights are recorded, the sampling is without replacement to increase data variability. In case that less nights are recorded, sampling is conducted with replacement. To account for possible duplicates, the values of the dependent variable, SH, and weight are distorted by uniform Gaussian noise. This is a standard data augmentation technique to produce synthetic data points (Beinecke and Heider, 2021). Further, this augmentation is known to increase the generalization properties and thus the explanatory power of a model (Shorten and Khoshgoftaar, 2019).

As a measure of feature importance, the permutation-based feature importance was chosen because it seems most suitable on the given dataset (Breiman, 2001). Every feature is assigned a value between 0% and 100% such that the values sum up to 100% representing the importance -higher values imply higher importance. The permutation-based importance of a feature is calculated as the increase in the model’s prediction error after permuting the feature’s values. If the increase is large, the feature is important because the model’s performance relies on the feature. In contrast, if shuffling the values does not change the model’s error, the feature is said to be unimportant. When several features measure similar effects, i.e., are highly correlated, feature importance is particularly meaningful when the importance of a “cluster” of similar features is defined as the sum of the importance scores of the contained individual features and the importance of the cluster is considered the relevant variable. Ward’s clustering algorithm is applied to the similarity matrix of the features to identify these clusters. There is no universal threshold above which a cluster is considered relevant; in particular, such a threshold must depend on the number of features. Indeed, given completely random data with p independently sampled features and independently generated labels, every feature will form its own cluster. Further, any cluster is equally unimportant such that the cluster importance scores will turn out to be approximately 1/p each. In the present study, 8 clusters are identified such that 1/p ∼ 12.5%. Therefore, only clusters that exceed this threshold are considered important.

It is well known that the feature importance can be misleading if strongly correlated features are present (Breiman, 2001). To determine the feature importance reliably and, primarily, to increase model performance, different pre-processing methods are well studied (Genuer et al., 2010; Gregorutti et al., 2017). One popular approach is to “decorrelate” the data. The goal of the decorrelation step is to turn the features into independent features. Intuitively, the part of a feature that can be explained by combining the other features is subtracted and the left-over is seen as the actual contribution of the feature. This is sometimes called residual learning in the machine learning context (Zhang and Chan, 2003; Dezfouli et al., 2019; Verdinelli and Wasserman, 2021). In the current study, the Gram-Schmidt decorrelation method was used to decorrelate the dataset which was established to decorrelate sets of features rather recently (Hyvärinen and Oja, 1997; Zhang and Chan, 2003). In this method, the residual expresses the part of a feature that cannot be explained by a linear combination of the covariances of the other features. A detailed comparison of this decorrelation-based feature importance with other notions of feature importance is given by Gerstorfer et al. (2023).

Beside requiring a measure for feature importance, in a second step it becomes crucial to not only determine notable features but to explain their influence on the nocturnal behavior. Recently, SHAP values were discovered as a method in machine learning to identify the influence of features’ values on the dependent variable (Lundberg and Lee, 2017). Those values are based on Shapley values, an old concept in mathematical game theory (Shapley, 1988). Formally speaking, the SHAP value is computed as the mean marginal contribution of each feature value across all possible values in the feature space. It is important to notice that the SHAP approach describes the effects of the model and not trends in the real raw data. This is good to identify effects that are not directly visible in the data but hidden in a complex structure which can be learned and used by the model. But this approach clearly only yields to sane descriptions if the model’s performance is decent. SHAP values and their application in data scientific models are recently well-studied and used in life science studies (Lundberg et al., 2018). A detailed and formal introduction is given by Lundberg and Lee (2017). In the current contribution, we visualize the SHAP values as beeswarm plots which we call SHAP plots. A SHAP plot contains three dimensions of information given by the x-axis, y-axis, and a color gradient. On the y-axis, the different features are given. If one point of the dataset is drawn, it contains exactly one value per feature. The value per feature is encoded by a color gradient from small to large. In the case of non-ordinal features, different colors represent different classes, but the gradient has no deeper meaning. The x-position of a point corresponds to the point’s SHAP value. Given a SHAP plot, trends and effects of features can be found by the means of finding clusters of colored points on the rows of the SHAP plot. For instance, if higher values of a feature correspond to lower SHAP values, the model suggests a negative trend and if smaller values imply a lower SHAP value, the model suggests a positive trend. But also, if there is a well-separated clustering of colors which is non-monotone, this corresponds to effects of the feature. If multiple separated clusters of the same color appear, effects cannot be described by the feature itself but also depend on other feature’s values.

## 3 Results

In the current study, three sets of results were obtained. These are presented in the following subsections. The first subsection provides a basic description of the nocturnal behavior of ungulates. In the next subsection, a model of nocturnal behavior was constructed and used to identify the most influential features. Furthermore, a SHAP analysis was conducted to identify the type of influence of each feature within the model. In the last subsection, the trends of the features found important by the model were examined on the actual dataset.

### 3.1 Description of nocturnal behavior

The activity budgets of the studied individuals are given in Figure 2. The three zebra species clearly differ from all other species. They show a lower proportion of lying (21.4% to 32.2%). Although the other species show a rather uniform picture with respect to the distribution of lying and standing, it is noticeable that the species of the genus *Tragelaphus* (67.8% in mean) as well as the Okapi (60.0%) show a slightly lower proportion of lying than the other Cetartiodactyla (79.6% in mean), while the genus *Hippotragus* have a higher proportion of lying (88.0% in mean). With respect to the proportion of LHD, the White Rhino shows the highest value (18.3%). On contrast, the Mountain Zebra (1.5%) and the Grevy’s Zebra (3.2%) have the lowest proportions of LHD whereas the proportion of LHD between all other species is comparable (min. 5% to max. 12.4%, in mean 9%).

**Figure 2:**
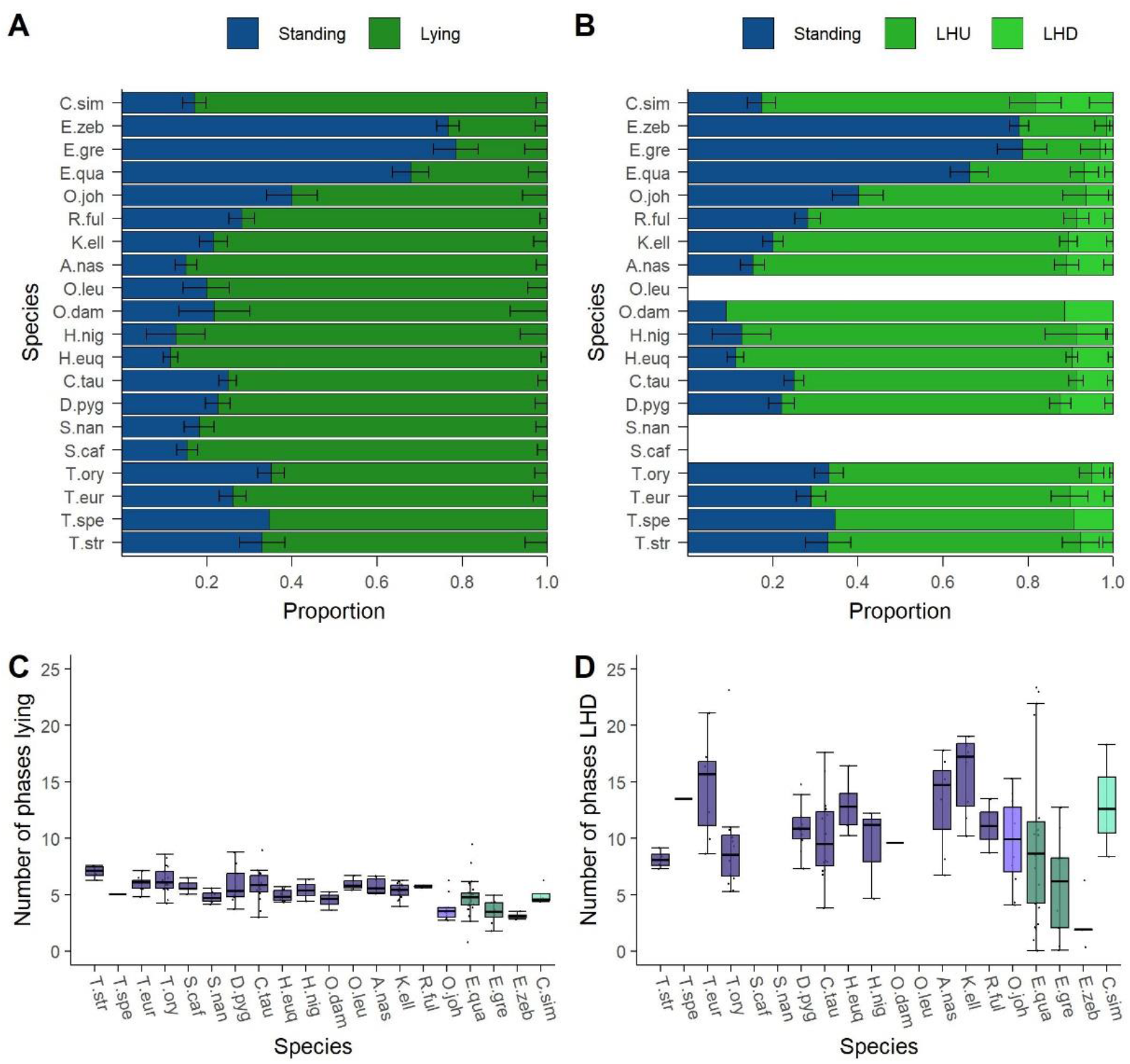
Description of the nocturnal behavior of the analyzed species’ adult individuals. The activity budget shows the proportion of the behaviors (A) standing and lying and (B) with lying split up into lying – head up (LHU) and lying – head down (LHD). The underlying data is normalized to the behavioral states excluding Out. The number of phases (C) lying and (D) lying – head down (LHD) are given in the second row of the figure. The families (Bovidae, Giraffidae, Equidae and Rhinocerotidae) are highlighted.

The number of phases of lying and LHD per night are visualized in Figure 2 (C) and (D). All species similarly often lie down per night with a mean of 5 phases, ranging from 3 to 7 phases. Noticeable, the smallest number of phases is shown by the Mountain Zebras (3 phases) and the Grevy’s Zebras (3.5 phases) as well as the Okapis (4 phases). On contrast, the largest number of lying phases is found within the genus *Tragelaphus* except the Sitatunga (6 to 7 phases). The number of phases LHD is much more variable between species and varies from 3 to 16 phases. In accordance with the proportion, the Mountain Zebras (3.4 phases) and the Grevy’s Zebras (4.7 phases) show the fewest phases of LHD, while the other species exhibit at least 7 such phases.

### 3.2 A model of nocturnal behavior

Based on above’s observation that the zebra’s behavior differs strongly from those of the Cetartiodactyla, we decided to model the influencing factors on nocturnal behavior only for the observed Cetartiodactyla to detect subtle effects. The nocturnal behavior is modelled by means of a random forest regression. Some of the analyzed features are strongly correlated and Ward’s clustering algorithm determines the clusters “age”, “sex”, “stable”, “month”, “zoo”, “taxonomy” (consisting of “genus” and “family”), “natural” (consisting of “food” and “habitat”), and “size” (containing “weight” and “SH”). The random forest models perform well as indicated by the explained variance scores R^2^ of 0.5 (proportion lying), 0.45 (proportion LHD), 0.29 (number of phases lying), and 0.55 (number of phases LHD).

The importance of any feature is given in Figure 3, for more detailed information, please refer to the table of values in the supplementary material (S5 Table). The individual’s age is highly relevant for all behavioral parameters. Furthermore, the cluster “taxonomy” is considered relevant for proportion lying and the number of phases LHD. In both cases, the relevance is due to the family while the genus cannot be considered important. Moreover, the animal’s size is relevant regarding the number of phases lying and LHD. Finally, the cluster “natural” is considered important with respect to the proportion of lying. Here, the dietary type (“food”) shows a larger relevance. Remarkably, the housing conditions (“stable” and the non-measurable factors “zoo”), the sex and the cluster “stable” do not show any relevance in the model.

**Figure 3:**
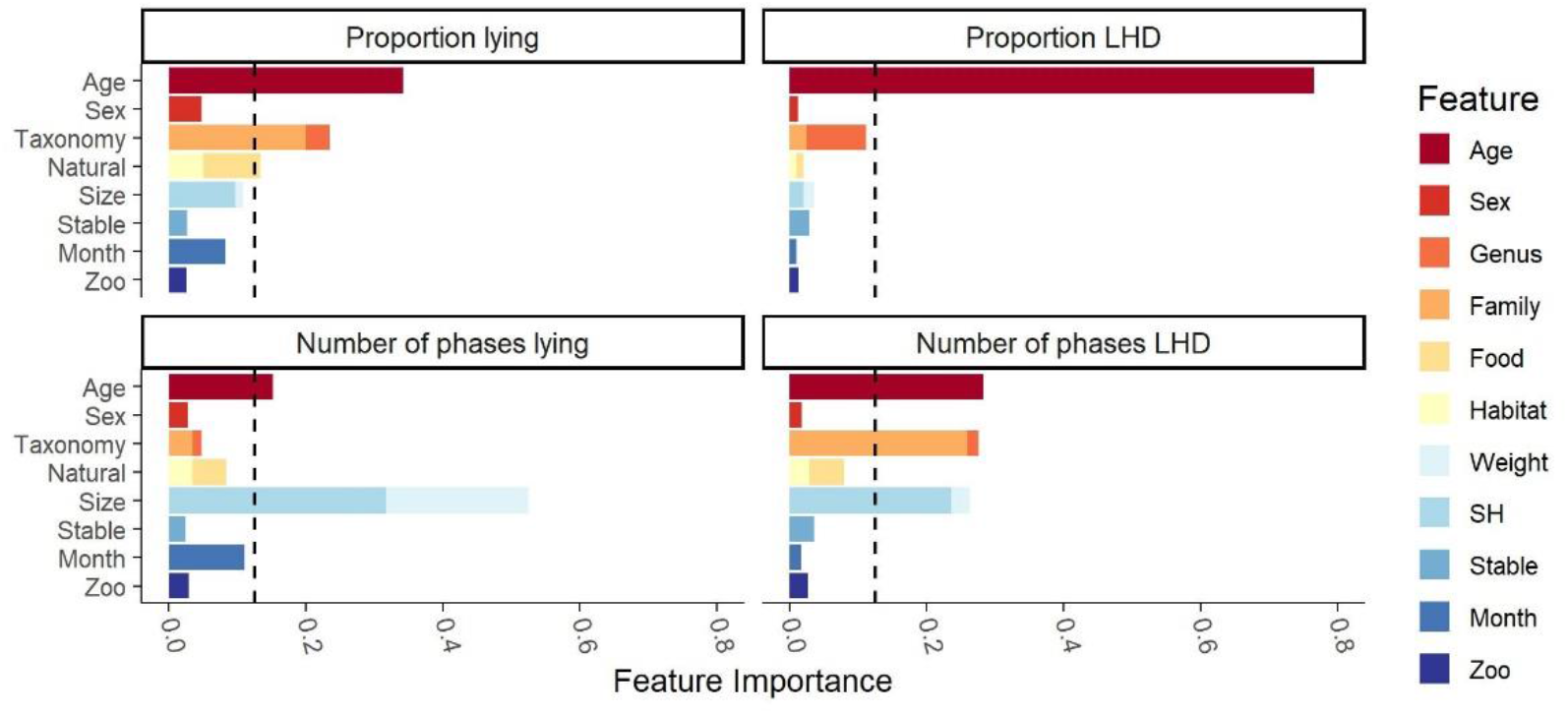
Feature importance of the clusters found by Ward’s clustering algorithm on the Cetartiodactyla’s data. Ward’s clustering algorithm identified the eight clusters (age, sex, taxonomy, natural, size, stable, month, zoo) given on the y-axis. The feature importance scores were measured by random forest regression with the Gram-Schmidt decorrelation method. The x-axis reports the feature importance score of any feature. The dashed line marks the threshold of 12.5% from which on a cluster was considered influential.

Beside determination of the features’ importance, the model can be used to determine effects of the single features on the model’s prediction. Figure 4 contains the SHAP plots for all four behavioral parameters. In the following, only the effects of previously found relevant clusters are presented. Regarding the individual’s age, the model predicts that juvenile individuals have higher proportions of LHD and lying and show more phases of LHD than adult and subadult individuals. A clear difference between subadult and adult individuals is only predicted for proportion lying: subadult individuals lay more than adult individuals. Regarding the taxonomy, a clear distinction between Giraffidae and Bovidae is predicted for the proportion and the number of phases lying. On a more fine-grained level, the values of the genus *Tragelaphus* are apart from the other Bovidae’s values, animals of the genus *Tragelaphus* show a smaller proportion of lying but a higher number of lying phases. With respect to the “natural” cluster, a distinction between the different dietary types becomes explicitly visible for the parameters regarding lying. Browsers are predicted to have more lying phases. Also, there is a small trend visible which indicates that grazers show a larger proportion lying than browsers. Regarding the cluster “size”, the model predicts an almost monotone decrease in the proportion LHD and the number of phases LHD with increasing size. This effect is visible with respect to the number of phases LHD for SH and weight while it is only visible for SH with respect to the proportion LHD. Moreover, a small increase in the proportion lying as well as the number of phases lying is predicted with increasing SH.

**Figure 4:**
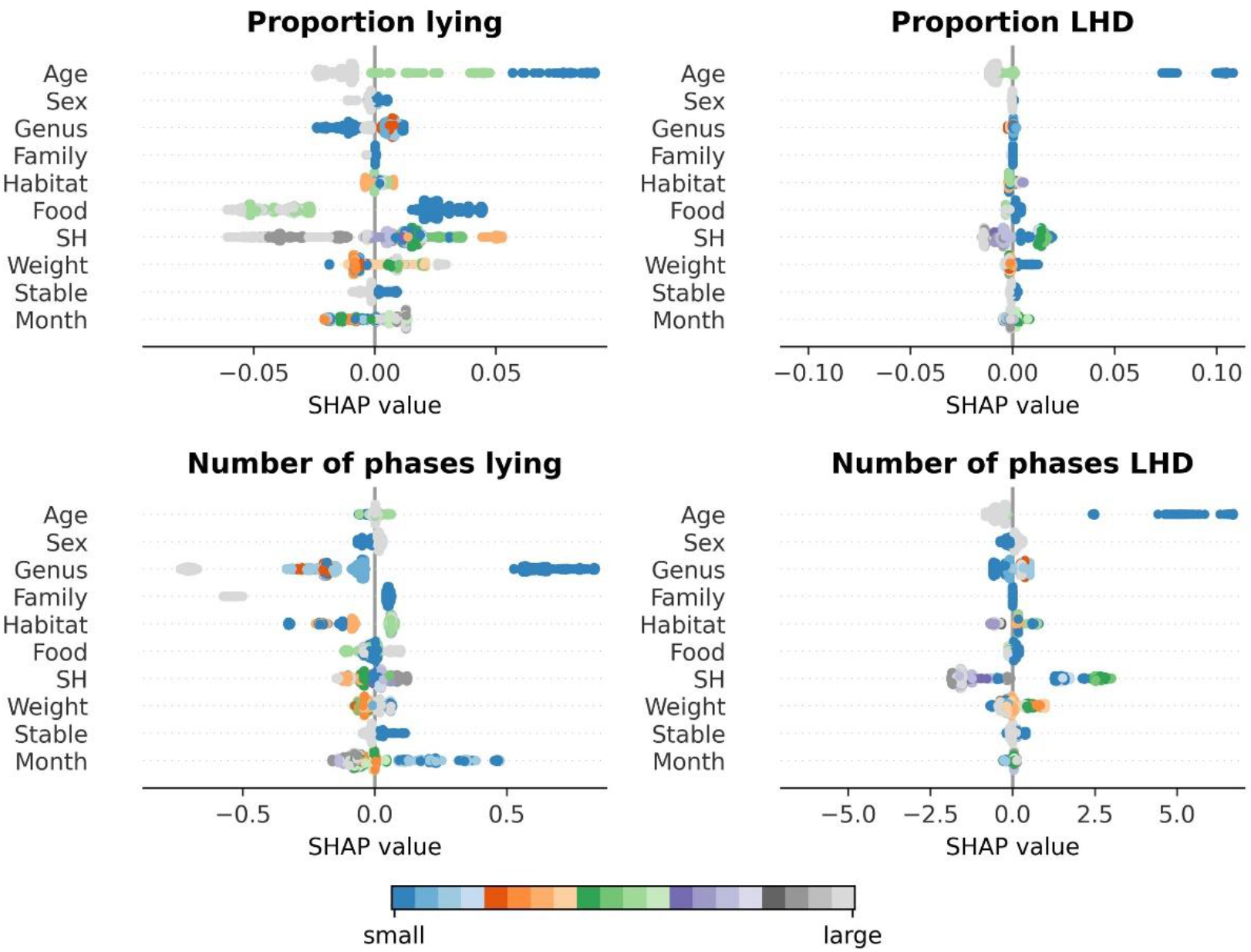
Trends as examined by a SHAP analysis of the random forest-based model on the Cetartiodactyla’s data. The feature’s value is indicated by the color gradient from small (blue) to large (grey). On the x-axis, the SHAP value is given. A positive SHAP value indicates that the feature’s value has a positive influence on the model’s prediction, a negative SHAP value indicates that the model’s prediction is reduced. The y-axis reports the features.

### 3.3 Examination of trends

In this subsection, those clusters identified to be important by the model, are analyzed in detail on the raw data. In contrast to the previous section, the full dataset is used, and the analysis is independent of the model.

Age is found to have a large impact on all behavioral parameters. The according trends are given in Figure 5 for those observed species for which data of multiple adult and non-adult individuals is present. With increasing age, the proportion of lying, proportion of LHD, and the number of phases LHD decreases. Regarding the cluster “taxonomy”, trends are already described in subsection “Description of nocturnal behavior”.

**Figure 5:**
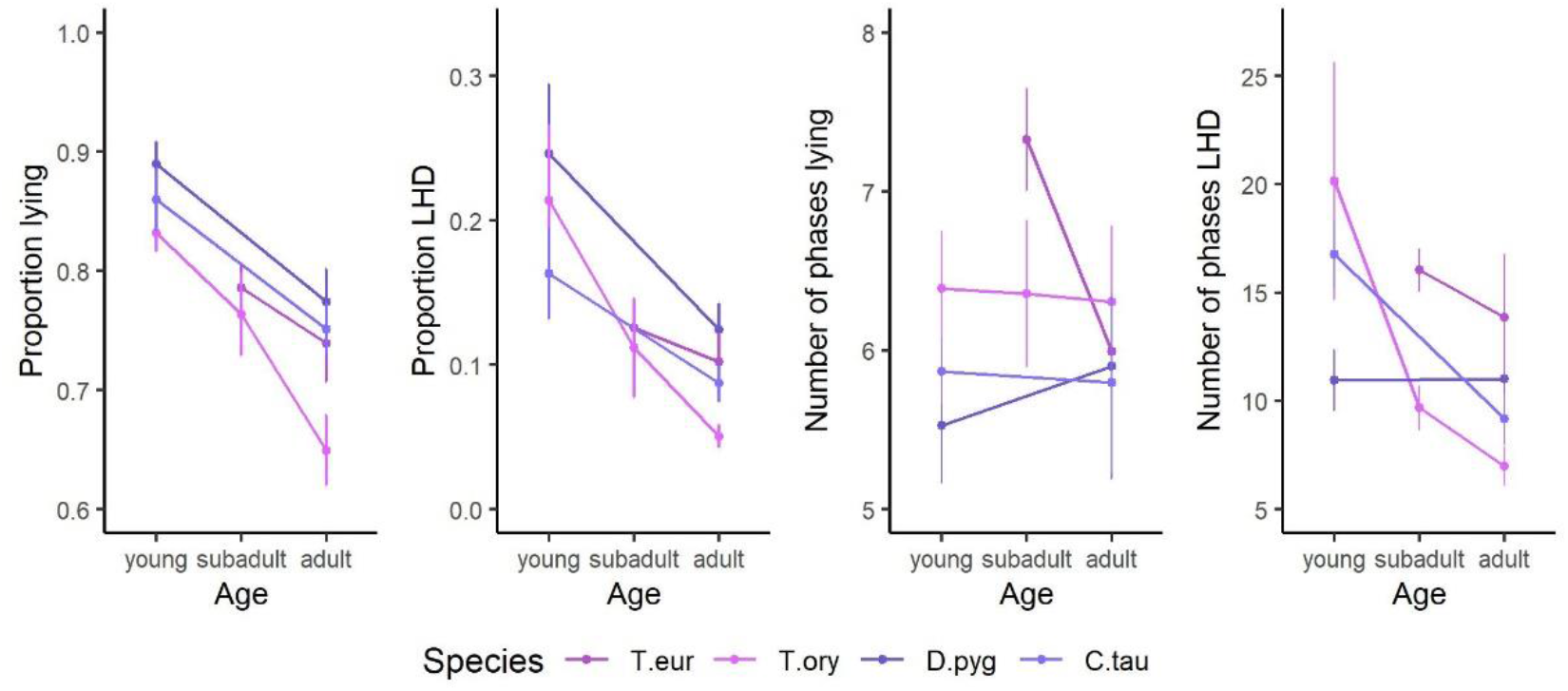
The trends of the influencing factor “age” on the features proportion lying, proportion lying – head down (LHD), number of phases lying, and number of phases lying – head down (LHD). The species’ mean and the SEM are given. Only species of the order Cetartiodactyla with at least two age categories and at least two individuals per age category are shown: Bongo (*Tragelaphus eurycerus)*, Common Eland (*Tragelaphus oryx*), Blesbok (*Damaliscus pygargus*), and Common Wildebeest (*Connochaetes taurinus*).

The feature “food” distinguishes between browsers, intermediate, and grazers. Figure 6 visualizes the differences between the different digestion types. As the family was found to have a large influence on the behavior, the Okapis are marked accordingly. Within the other species, browsers show a little bit smaller proportion of lying than grazers but more lying phases. Similarly, the same difference regarding the proportion LHD between browsers and grazers is indicated by the data.

**Figure 6:**
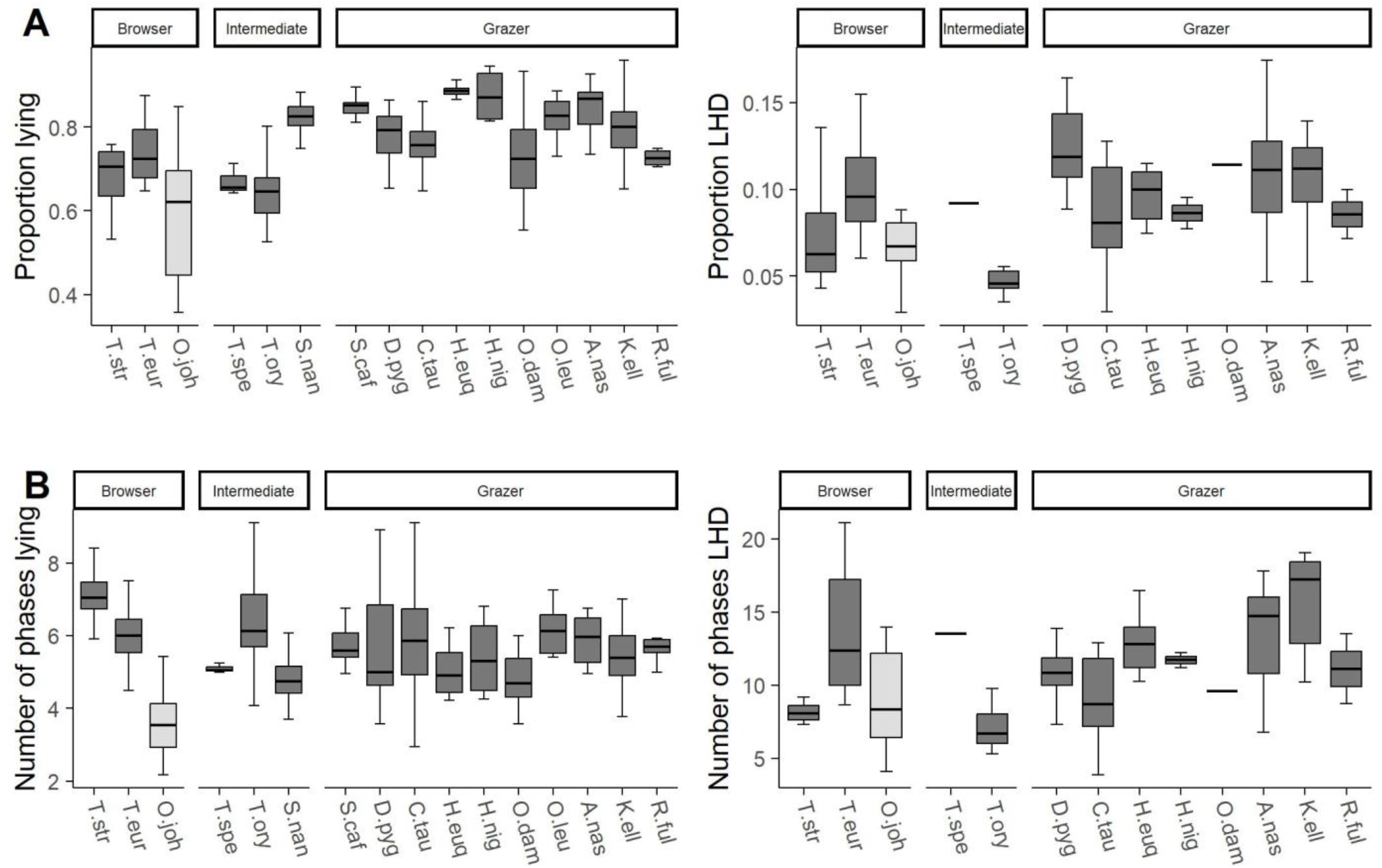
The trends of the influencing factor “food” on the features (A) proportion lying and proportion lying – head down (LHD) as well as (B) number of phases lying and number of phases lying – head down (LHD). On the x-axes, the species are given, and the three possible values of the feature food (browser, intermediate, grazer) are reported.

Finally, features of the cluster “size” shows a notable influence on the nocturnal behavior. The influence of the animal’s weight and its SH were analyzed in detail by linear regression models. As most behavioral studies consider the weight rather than SH, a visualization for the feature weight is given in Figure 7. There is a moderate negative correlation between the animal’s size and both, proportion LHD and the number of phases LHD, i.e., the larger a species, the less time it spends on LHD. For completeness, all correlation coefficients with respect to SH are negative and the coefficients of determination R^2^ read <0.01 (proportion lying), 0.31 (proportion LHD), <0.01 (number of phases lying), and 0.17 (number of phases LHD).

**Figure 7:**
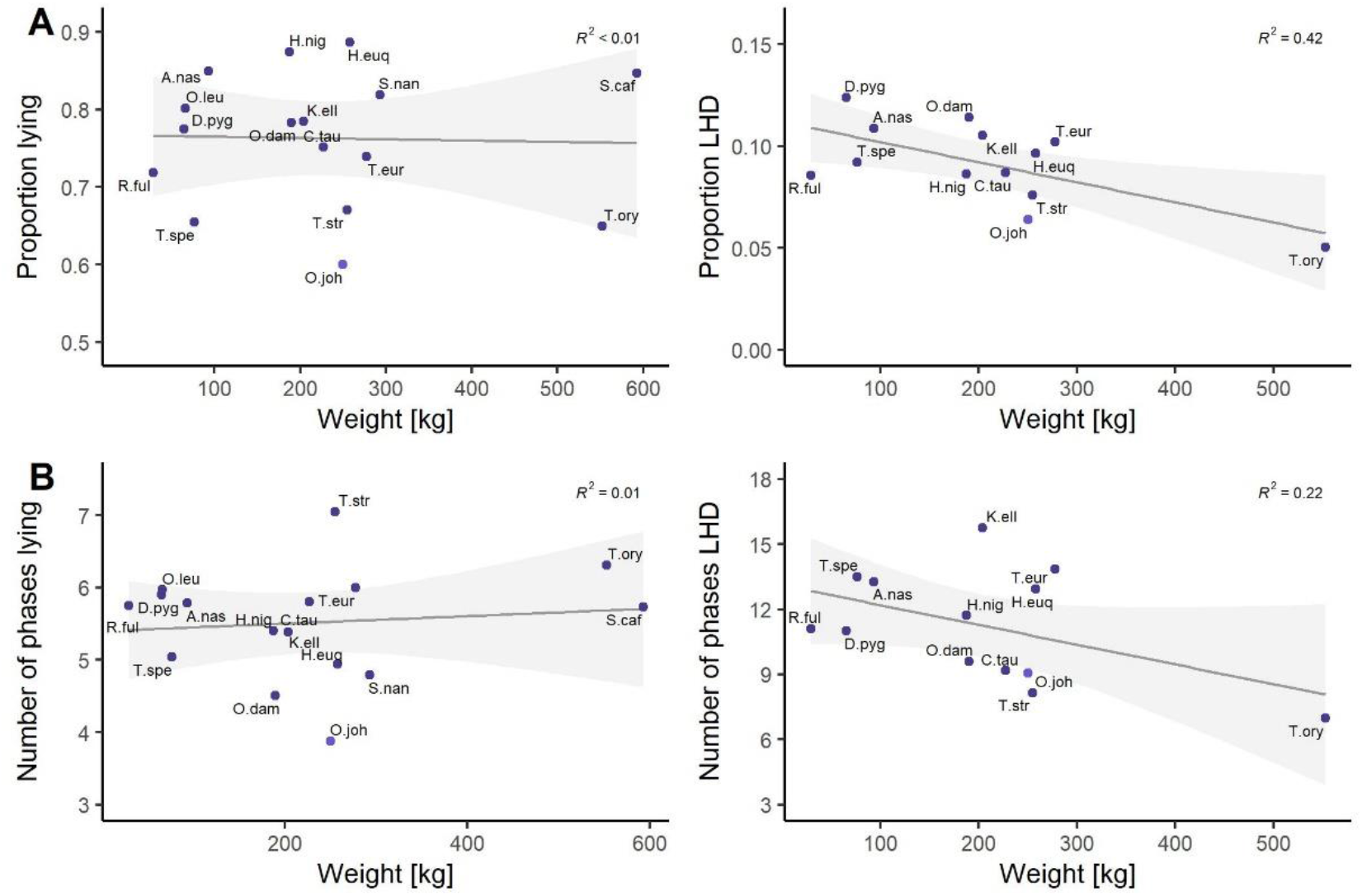
Linear regression models between the species’ weight and (A) the proportion lying and proportion lying – head down (LHD) and (B) the number of phases lying and number of phases lying – head down (LHD). The coefficient of determination is reported on the top-right in every subfigure. The families Giraffidae and Bovidae are highlighted.

## 4 Discussion

In this contribution the nocturnal behavior of 19 ungulate species was studied. The study contains, for some observed species, the first description of nocturnal behavior that we know of, both in the wild and in zoos. While the number of individuals per species varies (see S1 Table), the study was based on the evaluation of 196 individuals recorded over 47 nights on average and thus provides a good description of ungulates’ nocturnal behavior.

### 4.1 Description of nocturnal behavior

Regarding the general description of nocturnal behavior, existing studies are partly hard to compare to the current study (Santymire et al., 2012; Owen-Smith and Goodall, 2014; Davimes et al., 2016). First, if sleep is measured physiologically, some studies do not distinguish REM sleep and non-REM sleep, but they report the total sleep time TST which cannot be estimated by video recordings. Second, some behavioral studies define sleep as lying with closed eyes and do not refer to the typical REM sleep position. Third, even the comparison of values for lying is hard between some studies as they distinguish standing, lying, and ruminating and the latter can be done while standing or lying such that the time lying cannot be inferred.

On the presented data, a striking difference was observed between the three zebra species on the one hand and the members of the order Cetartiodactyla on the other hand (Figure 2 (A, B)). Ruckebusch (1972) examined the nocturnal behavior of farm animals including, among others, horses, and cows. His evaluation of standing and lying fits well into our findings. Horses lie 20% of the night and the three zebra species lie 25.6% on average. Cows were observed to lie 87.5% during night and, in the current study, Bovidae lie up to 88.6% of the time. Similarly, Ruckebusch (1972) found cows to be 6.3% of their time in the REM sleep position while Bovidae’s values in the current study range from 5.0% to 12.4% and the number of phases of REM sleep (0.83 per hour) compare well to the median values in the current study (0.64 to 1.4 phases per hour). Only the three observed zebra species spend a bit less time in the REM sleep position (1.5% to 6.8%) than the horses (7.8%) but the typical number of such phases of horses per hour (0.9) is comparable to the one of the three zebra species (0.3 to 0.9). The observation that zebras spend a lot more time standing and less time lying than the observed Cetartiodactyla is not surprising if the digestion type is considered as an explaining factor. Zebras, as hind-gut fermenters, need to devote more time to feeding and foraging due to a need for greater food intake than ruminants (Owen-Smith and Goodall, 2014). The studied Cetartiodactyla, as ruminants, spend more time resting as ruminating occurs a lot while lying down (Janis, 1976).

A recent study of two male free-roaming blue wildebeest was conducted by Malungo et al. (2021). The wildebeest were found to be 1.6% of the time in REM sleep divided up into 0.6 ± 0.04 phases per hour. Although the sample contains only two individuals, the values are in accordance with the current findings based on twelve adult individuals (2.9% to 12.8% and 0.35 to 1.17 phases per hour). Similarly, Davimes et al. (2018) found one Arabian Oryx to be 1.5% ± 2.5% time being in REM sleep with about 0.54 ± 0.15 phases per hour which compares well to the observations of Bovidae’s behavior in the present study. To summarize, findings of other studies regarding ungulates fit well into our description of nocturnal behavior and therefore, the presented data can be seen as a valid reference for further studies on ungulates.

### 4.2 Influencing factors on Cetartiodactyla’s nocturnal behavior

The random forest model performs well on the given dataset. The explained variance scores clearly exceed the threshold for a strong relation between the model’s predictions and the raw data (Cohen, 1988). Moreover, there is a very good fit between the trends on the relevant features in the model which were measured by the SHAP analysis, and the trends described on the raw data. In conclusion, the model seems to adequately describe the nocturnal behavior of the analyzed Cetartiodactyla and the classification of the relevant factors influencing the nocturnal behavior can be considered reliable.

The individual’s age was found to be the most dominating influencing factor of nocturnal behavior. Particularly, young individuals spend much more time lying and in the REM sleep position than adult animals (see Figure 5). This is not surprising as age has been proven to influence the activity/rest cycles of different mammals (Siegel, 2005; Ruckstuhl and Neuhaus, 2009), and age is known to be an influencing factor of REM sleep in mammals and birds (Ruckstuhl and Kokko, 2002; Cajochen et al., 2006; Steinmeyer et al., 2010; Rattenborg et al., 2017; Burger et al., 2021).

Furthermore, the animal’s size was identified as an important influencing factor. We decided to focus on the animal’s weight as this is a well analyzed factor in literature (Allison and Cicchetti, 1976; Siegel, 2005; Lesku et al., 2008; Gravett et al., 2017). It is known that there is a strong negative correlation between a species’ average weight and its REM sleep time. This correlation was observed on a very macroscopic level, from mice to African elephants (Siegel, 2005). On this scale, it is not possible to draw conclusions for species that are comparable in weight. However, the current study observes this effect on a very microscopic level. In particular, the more closely related analyzed species with comparable sizes show the same negative correlations.

### 4.3 Conclusion

The present study is a large-scale study on nocturnal behavior of various ungulates, more precisely it describes the basic behaviors standing, lying and being in the REM sleep position. To our knowledge, it provides the first description of nocturnal behavior for some species and extends the database for the other species significantly. The results fit well into the sparse existing literature. Thus, the data can be considered a valid reference for further research. Such data is not only relevant for scientific purposes but can also help to assess animal’s welfare in zoos. The most important influencing factors on nocturnal behavior were determined as the individual’s age, the species’ body measurements, the taxonomic relation as well as the feeding type.

## 6 Author Contributions

**JG:** Conceptualization (lead); Data curation (lead); Formal analysis (equal); Investigation (lead); Methodology (equal); Visualization (equal); Writing – original draft (lead). **MH:** Data curation (supporting); Formal analysis (equal); Methodology (equal); Visualization (equal); Writing – original draft (supporting). **PD:** Conceptualization (supporting); Funding acquisition (lead); Project administration (lead); Supervision (lead); Writing – original draft (supporting). All authors approved the submitted version.

## 7 Acknowledgments

The research leading to this publication was funded by von Opel Hessische Zoostiftung.

The authors gained great support from directors, curators, and animal keepers of the participating zoos (in alphabetical order of town): Königlicher Burgers Zoo Arnheim, Zoologische Gärten Berlin (Tierpark and Zoo), Zoo Vivarium Darmstadt, Zoo Dortmund, Zoo Duisburg, Zoo Frankfurt, Zoom Erlebniswelt Gelsenkirchen, Erlebnis-Zoo Hannover, Zoo Heidelberg, Kölner Zoo, Zoo Krefeld, Opel-Zoo Kronberg, Zoo Landau in der Pfalz, Zoo Leipzig, Allwetterzoo Münster, Zoo Neuwied, Zoo Osnabrück, Zoologischer Garten Schwerin, Der Grüne Zoo Wuppertal.

## Supplementary Material

### 9 S1 Table: Sample sizes of individuals and nights within the given dataset

Overview of the distribution of the given dataset. For every species, the number of evaluated nights per individual is given. The first column contains the name of the evaluated species. For any species, there are two rows. The first row contains the number of evaluated individuals per age category, in brackets it is given for how many individuals Lying could be distinguished into LHU and LHD. The second row contains the corresponding number of evaluated nights. Species in the last two rows are not used to build the random forest model. Overall, 196 individuals with 9,239 nights containing 101,629 hours of video material were analyzed. The minimum number of nights per one individual is 9 nights and the maximum is 185 nights. The median is 46.5 and the mean amounts 47.1. The corresponding quartiles are 34 and 55 nights. LHU and LHD were analyzed on 132 individuals with 6,265 nights containing 68,915 hours of video material.

### 10 S2 Table: Performance of the deep learning-based software package BOVIDS

This table gives an overview about the performance of the used deep learning-based software package BOVIDS. To guarantee that the obtained results are valid, a total of 517 nights consisting of 5,687 hours of video material were prepared manually as a testing set. The most important measures of performance are contained: the f-score for the detection of Lying and LHD by the system and a comparison of how well the behavioral parameters are predicted per tested individual and per species. Overall, an average f-score of 0.992 ± 0.003 (Lying) and 0.956 ± 0.006 (LHD) is achieved. Accordingly, proportion Lying is estimated correctly up to 0.001 and proportion LHD up to 0.002 on average. Finally, the average number of phases Lying is overestimated by 0.4% and the number of phases LHD by 5.7%. All data of the White Rhinos were evaluated manually.

**Figure S1:**
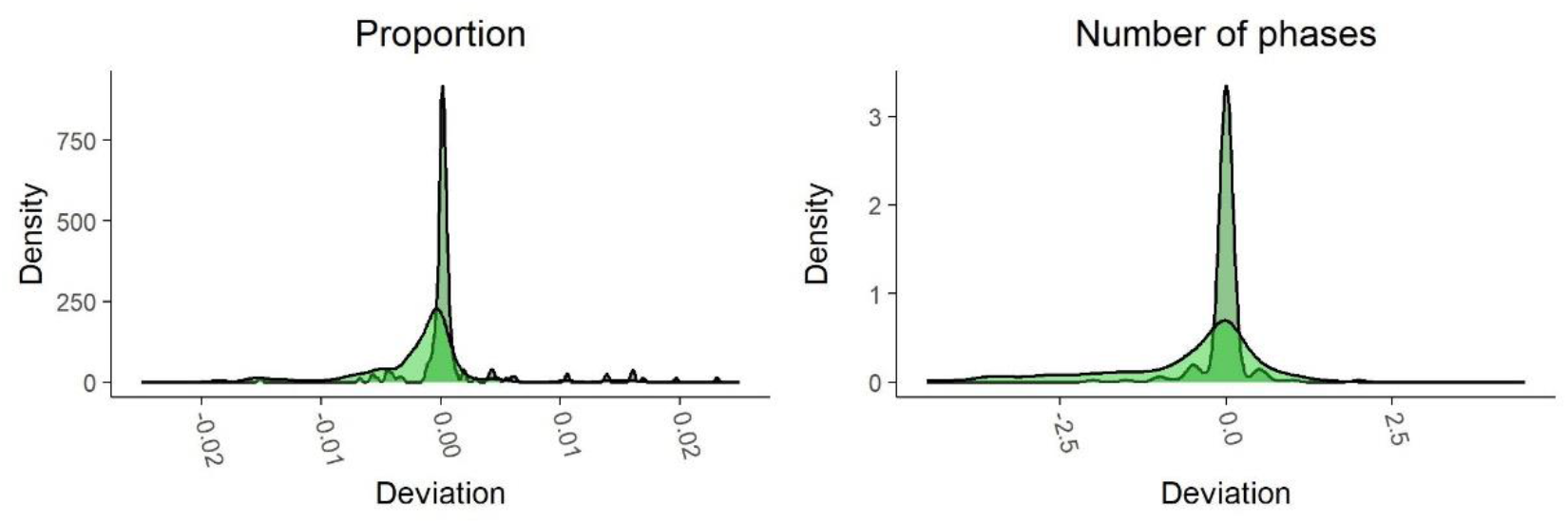
Visualization of the data shift between the manual annotation and the automatic evaluation by BOVIDS on the testing set. With respect to the proportion, outliers are dismissed in the graphic and can be found in S2 Table. The outliers are almost symmetric around zero. All distributions have a strong peak at zero which indicates that no severe systematic errors are produced.

### 11 S4 Table: Species describing influencing factors

Contains the information used in the modeling of influencing factors regarding the species. The systematic classification of the studied species is shown. Furthermore, the habitat is given a number to distinguish closed environments (3) from open environments (0) following the categorization of the IUCN Red List of Threatened Species (IUCN, 2022). For describing the body measurements shoulder height (SH), and weight is used. Also, the dietary type of browser, intermediate, and grazer is noticed. All information besides the habitat is gathered from different sources (Grzimek, 2000; Puschmann et al., 2009; Groves and Leslie Jr, 2011; Rubenstein, 2011; Skinner and Mitchell, 2011; Tacutu et al., 2013; Myers et al., 2021).

### 12 S5 Table: Data underlying the figures in the results’ section

Tabular representation of the values used for plotting Figure 2 and Figure 3.

